# Loss of stromal VEGFR2 protects against ischemic acute kidney injury by preserving the renal microvasculature

**DOI:** 10.64898/2026.10.27.762337

**Authors:** Takuto Chiba, Akira Oda, Ayako Kinoshita, Kaori Kamijo, Aryana Bhasin, Débora M. Cerqueira, Elina Mukherjee, Katherine E. Pfister, Yuko Hidaka, Yukiko Sakurahara, Katherine V. Maringer, Natasha M. Rogers, Jacqueline Ho, Sunder Sims-Lucas

**Affiliations:** Department of Pediatrics, University of Pittsburgh School of Medicine, UPMC Children’s Hospital of Pittsburgh, Pittsburgh, Pennsylvania; Department of Nephrology and Hypertension, Fukushima Medical University, Fukushima, Japan; Department of Pediatrics, Showa Medical University, Tokyo, Japan; Centre for Transplant and Renal Research, Westmead Institute for Medical Research; Department of Pediatrics, University of Maryland, Baltimore, School of Medicine

## Abstract

Injury to the renal microvasculature, particularly the peritubular capillaries, is a defining feature of ischemic acute kidney injury (AKI). Vascular endothelial growth factor receptor 2 (VEGFR2) signaling is a central regulator of endothelial cell survival, proliferation, migration, and vascular remodeling. Although VEGFR2 has been extensively studied in endothelial cells, accumulating evidence suggests that it is also expressed in non-endothelial cell populations, where it may regulate tissue-specific responses to injury. Here, using publicly available single-cell or single-nucleus RNA sequencing data from human and mouse kidneys, we identified VEGFR2 expression in renal stromal populations. Immunostaining further detected phosphorylated VEGFR2 in the renal stroma of mouse kidneys. To determine the functional role of stromal VEGFR2, we utilized renal stroma-specific *VegfR2* KO mice. Stromal *VegfR2* deletion attenuated kidney injury after renal ischemia/reperfusion injury (IRI), as evidenced by reduced tubular injury, preservation of renal blood flow, decreased endothelial oxidative stress, and maintenance of the renal microvasculature. Stromal *VegfR2*-deficient kidneys also exhibited increased *Angpt1* and *Tek* levels, enhanced endothelial ERK activation, and preserved endothelial adherent junctions. Taken together, these findings reveal an unexpected role for VEGFR2 in the renal stromal lineage and highlight stromal–endothelial communication as a potential therapeutic target for preserving microvascular integrity after ischemic AKI.

## Introduction

Injury to the renal microvasculature, particularly the peritubular capillaries, is a defining feature of ischemic AKI (1, 2). This injury is mediated, in part, by vascular oxidative stress. Although angiogenic responses are essential for endothelial repair, the renal microvasculature has a limited capacity for regeneration following injury(2). Therefore, therapies that promote renal microvascular repair represent a promising strategy to improve recovery from ischemic AKI and prevent progression to CKD.

Vascular endothelial growth factor A (VEGF-A) is a key regulator of endothelial cell function and vascular homeostasis. Binding of VEGF-A to its principal receptor, VEGF receptor 2 (VEGFR2), activates signaling pathways that promote endothelial cell survival, proliferation, migration, and vascular remodeling(3, 4). Although VEGFR2 has been extensively studied in endothelial cells, accumulating evidence suggests that it is also expressed in non-endothelial cell populations, where it may regulate tissue-specific responses to injury(5, 6). In the kidney, coordinated communication between renal stromal cells and endothelial cells is essential for maintaining microvascular integrity and supporting adaptive vascular repair following obstructive injury(7). Although VEGFR2 signaling is generally considered vasculoprotective, whether its activity in renal stromal cells promotes or limits microvascular preservation after ischemic AKI remains unknown.

Here, we investigated the role of VEGFR2 in the renal stromal lineage using a Foxd1-Cre-mediated cell type specific loss of function model. Foxd1-expressing renal stromal progenitors give rise to heterogeneous cell populations, including pericytes, vascular smooth muscle cells, mesangial cells, and interstitial fibroblasts. We demonstrate that VEGFR2 is expressed in renal stromal cell populations in both human and mouse kidneys and provide evidence of stromal VEGFR2 activation in mouse kidneys following ischemic AKI. Unexpectedly, loss of stromal VEGFR2 preserves renal perfusion and microvascular integrity, reduces endothelial oxidative stress, and is associated with increased expression of ANGPT1–Tie2 pathway components. These findings identify stromal VEGFR2 as a previously unrecognized regulator of the renal microvascular response to ischemic AKI.

## Results

### VEGFR2 is expressed and activated in renal stromal cells following kidney injury

We explored cell-type-specific expression of VEGFR2 (also known as KDR) using publicly available single-cell RNA sequencing (scRNA-seq) and single-nucleus RNA sequencing (snRNA-seq) data from human and mouse kidneys. In human AKI kidneys, VEGFR2 was found to be modestly expressed in the activated fibroblasts (aFIB) population (**Figure 1A**). The aFIB cluster comprises activated fibroblasts involved in adaptive repair, maladaptive remodeling, and tissue repair processes. Positive fold-change values indicate that KDR is expressed at higher levels in this cell cluster relative to other cell populations. Analysis of an independent mouse snRNA-seq dataset from kidneys 14 days after unilateral ureter obstruction (UUO) revealed moderate *Kdr* expression in activated fibroblast populations (**Figure 1B**). Although transcript abundance was relatively low, these findings provide evidence consistent with VEGFR2 expression in non-endothelial stromal populations, where its functional significance remains poorly understood. VEGFR2 is phosphorylated when activated. Our immunostaining revealed that phosphorylated VEGFR2 (phospho-VEGFR2, Tyr1175) is expressed in the renal stroma in uninjured kidneys and 7 days after renal ischemia reperfusion injury (IRI; **Figure 1C**).

**Figure 1.**
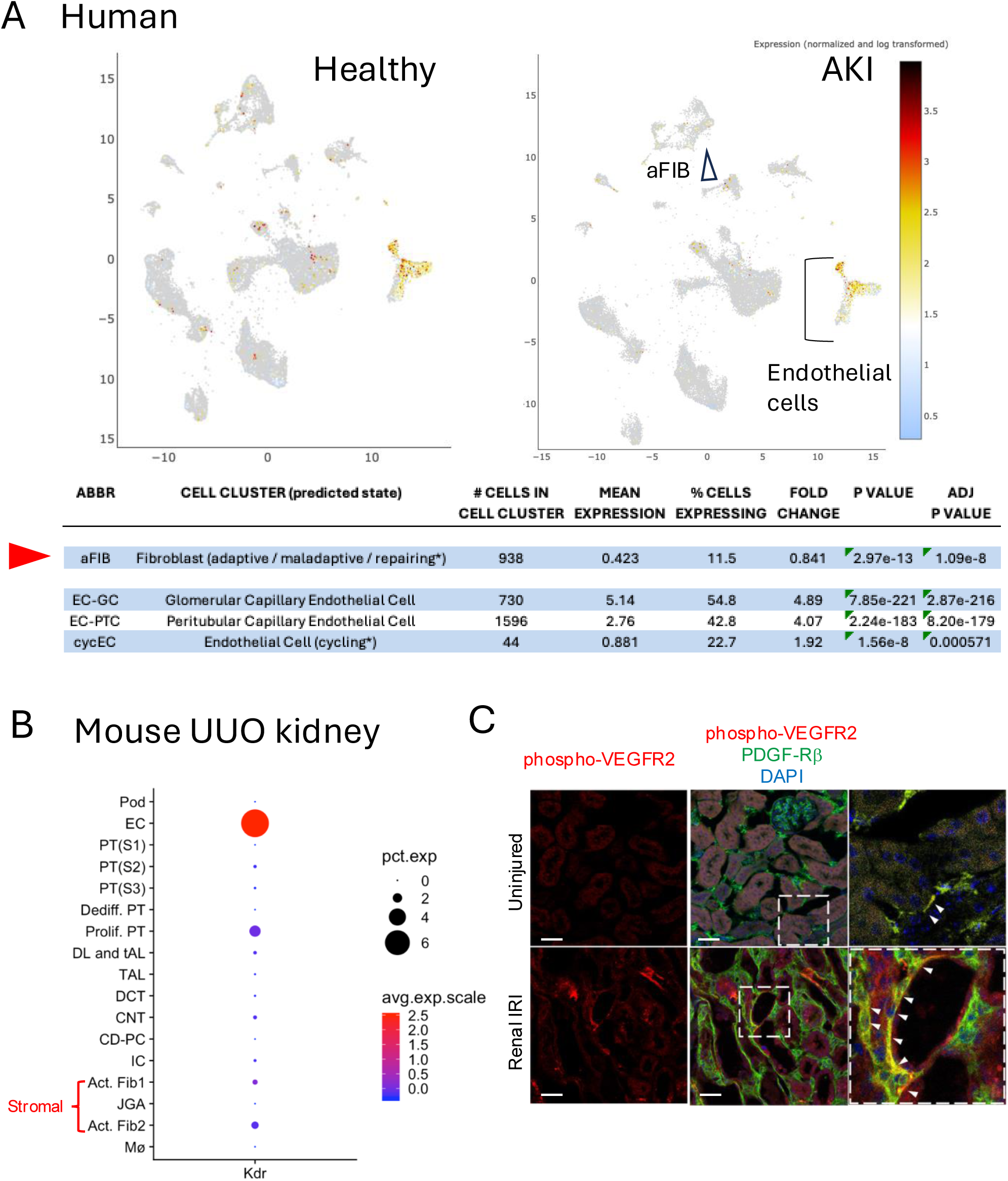
VEGFR2 is expressed and activated in renal stromal cells following kidney injury. **A.** Expression distributions of KDR (VEGFR2) in renal cell clusters from healthy and AKI human kidneys. Healthy, n = 28; AKI, n = 14. Data were derived from the Kidney Precision Medicine Project (KPMP) Kidney Tissue Atlas, which was accessed on August 20, 2025. **B.** *Kdr* expression was queried using the publicly available mouse kidney single-nucleus RNA-seq dataset from normal and day 14 unilateral ureteral obstruction (UUO) kidneys (Wu et al. PMID: 30510133) via the Humphreys’s laboratory KIT Kidney Interactive Transcriptomics website (http://humphreyslab.com/SingleCell/). **C**. VEGFR2 activation in renal stroma. Representative anti-phospho-VEGFR2 immunofluorescence staining in WT mouse kidneys under uninjured conditions and 7 days after renal IRI, co-stained with PDGFRβ.

### Stromal VEGFR2 deletion protects against ischemic AKI in mice

Previously, it was suggested that maladaptive cross-talk between VEGFR2 in endothelial cells and PDGFRβ in pericytes mediates kidney disease progression(7). However, the role of VEGFR2 in the renal stroma was unknown. To investigate the role of VEGFR2 in the renal stromal lineage, we used *Foxd1-Cre; VegfR2^fl/fl^*mice (hereafter *VegfR2^RSC−/−^*), in which *VegfR2* is deleted in Foxd1-lineage cells. Our laboratory previously characterized *VegfR2^RSC−/−^*mice, demonstrating efficient Foxd1-lineage-specific *VegfR2* deletion(6). We confirmed that no difference was noted in renal function and renal histology in the absence of injury between *VegfR2^RSC−/−^*and control kidneys in 10- to 14-week-old mice (**Figure 2A, D**, uninjured). *VegfR2^RSC−/−^*and control mice were next subjected to an ischemic AKI model in which we used unilateral renal IRI, followed by delayed contralateral nephrectomy 1 day before the samples were collected on day 7 (Figure 2). *VegfR2^RSC−/−^*male mice had decreased kidney injury 7 days after renal IRI, assessed by decreased serum levels of BUN and creatinine (**Figure 2B**). Further biochemical analysis revealed decreased phosphorus levels and increased Na/K and bicarbonate levels in *VegfR2^RSC−/−^* mice, suggesting renoprotective effects from the mutation (**Supplemental Figure 1**). Histology and immunostaining further confirmed reduced renal tubular injury in renal cortex area as well as reduced expression of kidney injury markers of KIM-1 and NGAL in *VegfR2^RSC−/−^* kidney tissue (**Figure 2C**). We also performed corresponding experiments in female mice and observed a similar protective phenotype following renal IRI, indicating that the renoprotective effects of stromal VEGFR2 deletion were observed in both male and female mice (**Figure 2D** and **E**).

**Figure 2.**
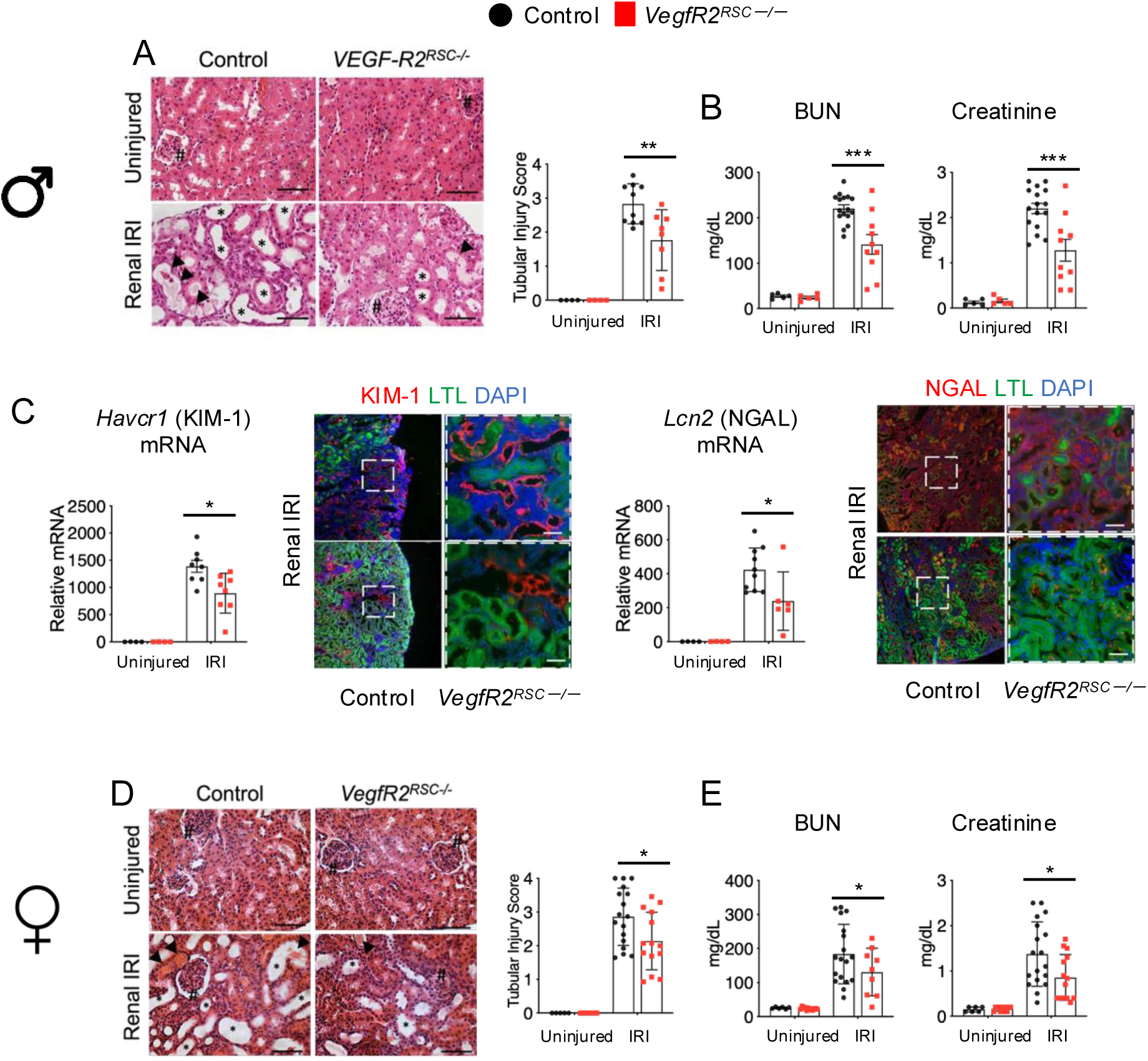
Stromal VEGFR2 deletion protects against ischemic AKI in both male and female mice. (A–C) Male mouse studies. (D and E) Female mouse studies. (A and D) Representative H&E-stained kidney sections and semiquantitative tubular injury scores demonstrate reduced renal tubular injury in *VegfR2^RSC^*^ー^*^/^*^ー^ mice 7 days after renal IRI. Glomeruli are indicated by #, dilated tubules by *, and proteinaceous casts by arrowheads. (B and E) Serum BUN and creatinine levels indicate reduced renal functional impairment in *VegfR2^RSC^*^ー^*^/^*^ー^ mice 7 days after renal IRI. (C) Renal *Havcr1* (KIM-1) and *Lcn2* (NGAL) mRNA levels and representative KIM-1 and NGAL immunofluorescence staining in male mice 7 days after renal IRI. Scale bars: 50 μm. n = 4–16. *P < 0.05, **P < 0.01, ***P < 0.001; one-way ANOVA followed by Tukey’s multiple-comparisons test.

### Stromal VEGFR2 deletion preserves renal blood flow after renal IRI

Ischemic AKI in mice is also characterized by reduction of renal blood flow (RBF) and microvascular rarefaction, which is often associated with incomplete endothelial repair after injury(2, 8, 9). We next hypothesized that *VegfR2^RSC–/–^* protects the function of the renal vasculature. Arterial spin labeling (ASL)-MRI allows noninvasive/longitudinal evaluation of RBF with its excellent quantitative capacity(10). Longitudinal ASL-MRI analysis demonstrated a significant genotype × time interaction (mixed-effects model, P = 0.0287), indicating that renal blood flow recovery differed between genotypes. *Post hoc* analysis showed significantly higher renal blood flow in the cortex area of *VegfR2^RSC–/–^* kidneys at day 7 after renal IRI (**Figure 3A**). Consistent with this, CD31-positive vascular area was preserved in *VegfR2^RSC–/–^* 7 days post renal IRI when compared with controls (**Figure 3B**).

**Figure 3.**
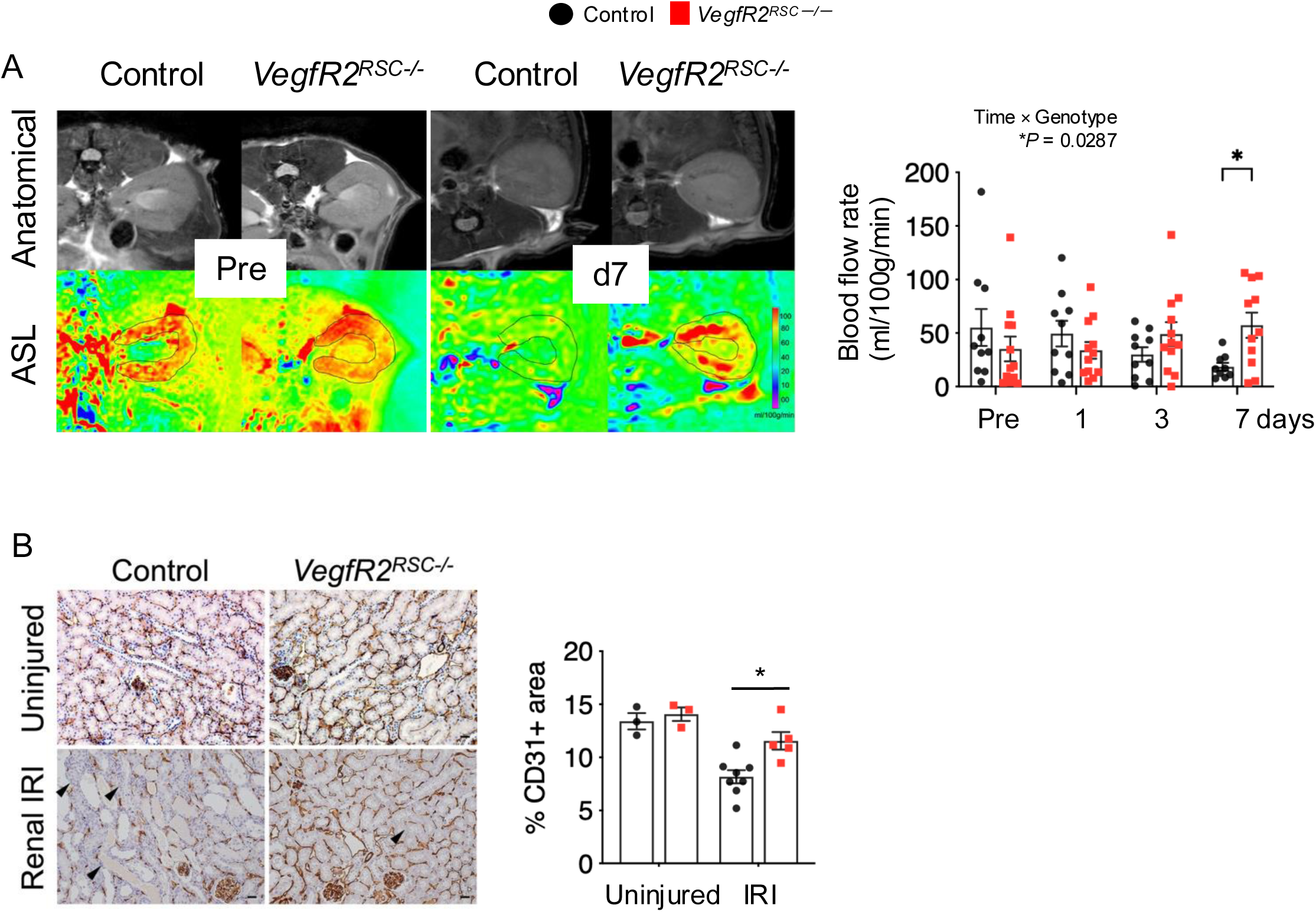
Stromal VEGFR2 deletion preserves renal blood flow after renal IRI. (**A**) Longitudinal renal blood flow was assessed by arterial spin labeling magnetic resonance imaging (ASL-MRI) before and at days 1, 3, and 7 after renal IRI. Representative anatomical and ASL-MRI images obtained before injury and at day 7 are shown. Renal cortical blood flow was quantified using ASL-MRI. A mixed-effects model (REML) revealed a significant genotype × time interaction (P = 0.0287), with significantly higher renal cortical blood flow in *VegfR2^RSC^*^ー^*^/^*^ー^ mice than in controls at day 7 after IRI, as determined by post hoc Tukey’s multiple-comparisons test. (**B**) CD31-labeled renal endothelial cells are increased in *VegfR2^RSC^*^ー^*^/^*^ー^ kidneys 7 days post renal IRI, whereas there is no change in uninjured kidneys. *p < 0.05; using 1-way ANOVA *post hoc Tukey* multiple comparison.

### Stromal VEGFR2 deletion reduces oxidative stress and vascular damage after renal IRI

Oxidative stress promotes endothelial cell senescence, which impairs angiogenesis and limits vascular repair after tissue injury(11, 12). Thus, we measured renal isofuran levels, a well-established *in vivo* oxidative stress marker(13). Renal isofuran levels were decreased in *VegfR2^RSC–/–^*kidneys 7 days post renal IRI (**Figure 4A**). Further, 4-hydroxynonenal (4-HNE) staining in renal endothelial cells was reduced in *VegfR2^RSC–/–^* kidneys 7 days after renal IRI (**Figure 4B**). Taken together, lack of VEGFR2 in the renal stroma decreases renal oxidative stress in the kidneys, especially in renal endothelial cells post renal IRI. Furthermore, the mRNA levels of genes associated with vascular injury, including *Syndecan-1* (*Sdc1*), *S100 calcium-binding protein A8* (*S100a8*), and Plasminogen Activator Inhibitor-1 (PAI-1, gene name *Serpine1*), were lower in *VegfR2^RSC–/–^* kidneys than in controls 7 days after renal IRI (**Figure 4C**). Immunostaining of PAI-1 showed consistent results that the number of PAI-1+ renal endothelial cells as well as renal interstitial cells was reduced in *VegfR2^RSC–/–^* kidneys (**Figure 4C** and **Supplementary Figure 2**).

**Figure 4.**
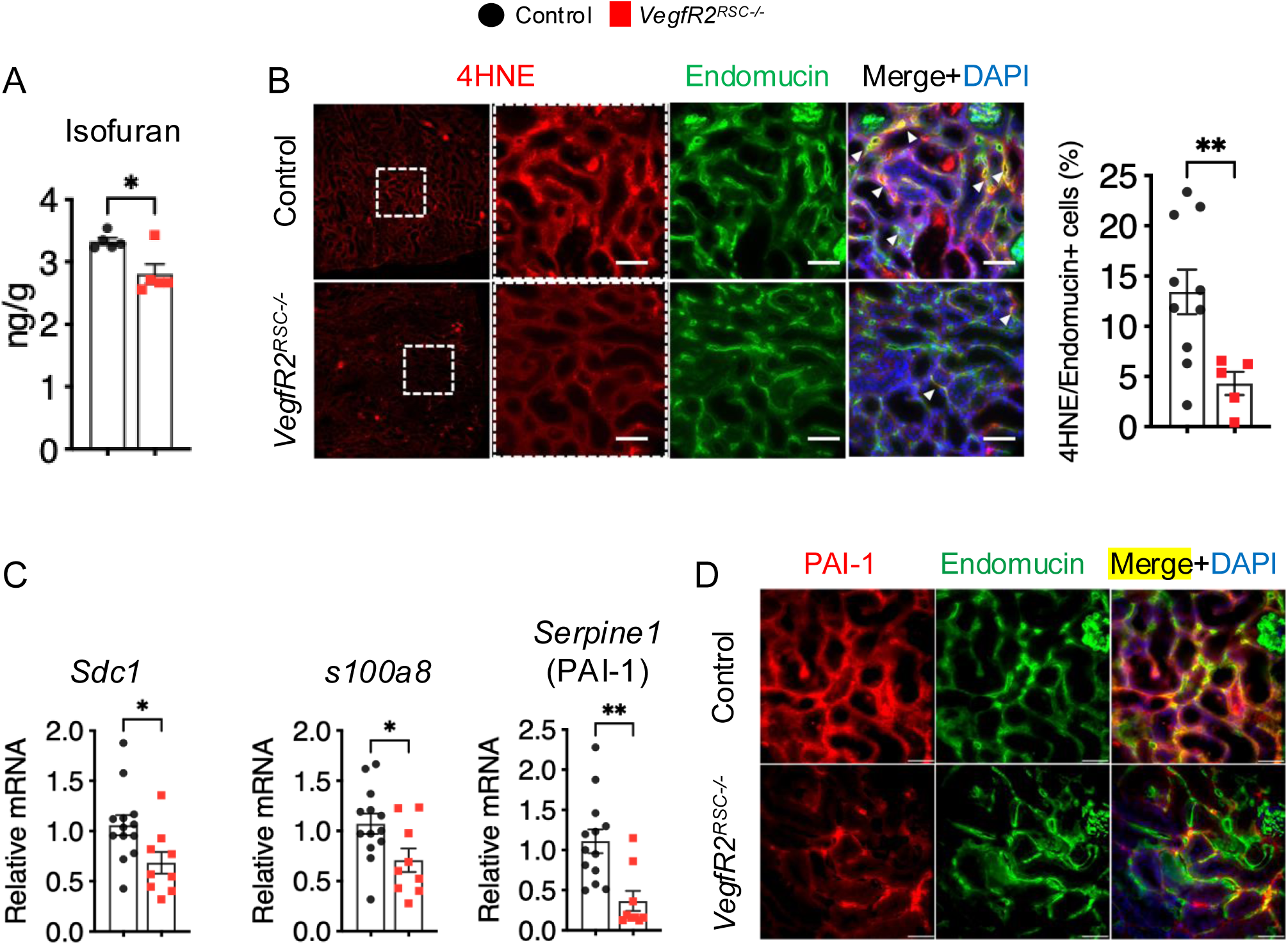
Stromal VEGFR2 deletion mitigates endothelial oxidative stress and vascular injury after renal IRI. (A) The *in vivo* oxidative stress marker, isofuran, is decreased in *VegfR2^RSC^*^ー^ *^/^*^ー^ kidneys 7 days after renal IRI. (B) Endomucin +/4-HNE+ oxidative stress-positive endothelial cells are reduced in *VegfR2^RSC^*^ー^*^/^*^ー^ kidneys 7 days after renal IRI. (C) *VegfR2^RSC^*^ー^*^/^*^ー^ kidneys decreased mRNA of vascular injury associated genes, *Sdc1, S100a8*, and *Serpine1.* Endomucin was co-stained as an endothelial marker. (D) *VegfR2^RSC^*^ー^*^/^*^ー^ kidneys decreased immunostained endothelial cells for PAI-1. Scale bars: 50 μm. *P < 0.05; **P < 0.01; ****P < 0.0001, using 1-way ANOVA *post hoc Tukey* multiple comparison or *t* test.

### Stromal VEGFR2 deletion is associated with enhanced endothelial stabilization after renal IRI

The ANGPT1–Tie2 signaling axis is a key regulator of endothelial survival, vascular stability, and angiogenic repair(3–5). We therefore examined whether components of this pathway were altered in *VegfR2^RSC–/–^*kidneys following ischemic AKI. We found that mRNA levels of *Angpt1* and *Tek* (Tie2) were higher in *VegfR2^RSC–/–^* kidneys than in controls 7 days after renal IRI (**Figure 5A**). Interestingly, ANGPT1 was expressed by PDGFRβ-positive renal stromal cells, although expression was also observed in tubular cells. Because phosphorylation of ERK is a marker of ERK activation, we next examined phospho-ERK expression. Phospho-ERK was increased in endothelial cells of *VegfR2^RSC–/–^* kidneys 7 days after renal IRI (**Figure 5B**). Furthermore, the expression of genes associated with vascular integrity, including *Esm1* (endothelial cell-specific molecule 1), *Proc* (protein C), and *Sdc2* (syndecan-2), increased in *VegfR2^RSC–/–^* kidneys 7 days after renal IRI (**Supplementary Figure 3**).

**Figure 5.**
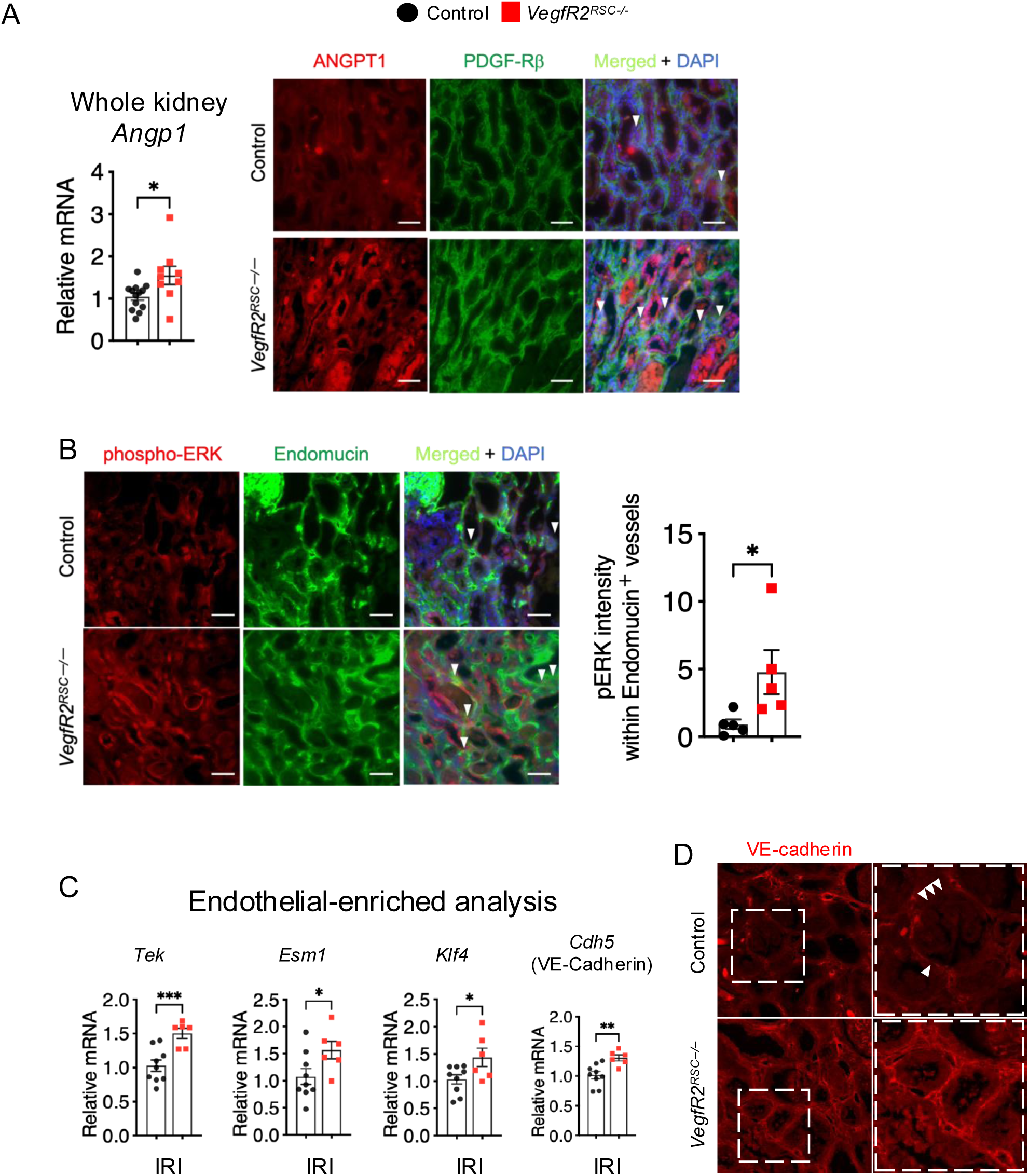
Stromal VEGFR2 deletion is associated with enhanced endothelial stabilization after renal IRI. (A) Whole-kidney *Angpt1* mRNA levels and representative ANGPT1 immunofluorescence staining with PDGFRβ in control and *VegfR2^RSC^*^ー^*^/^*^ー^ kidneys. (B) Representative phospho-ERK and Endomucin immunofluorescence staining and quantification of phospho-ERK intensity within Endomucin-positive vessels 7 days after renal IRI. (C) Relative mRNA levels of *Tek*, *Esm1*, *Klf4*, and *Cdh5* in CD31_⁺_-enriched renal cells isolated 7 days after IRI. (D) Representative VE-cadherin immunofluorescence staining demonstrating more continuous junctional staining in *VegfR2^RSC^*^ー^ *^/^*^ー^ kidneys compared with controls. Arrowheads indicate discontinuous VE-cadherin staining in control kidneys. Scale bars: 50 μm. *P < 0.05, **P < 0.01, ***P < 0.001; unpaired t test.

Finally, using CD31 -enriched renal cells from control or *VegfR2^RSC–/–^* kidneys 7 days after renal IRI, we found increased mRNA levels of *Tek, Esm1, Klf4*, and *Cdh5* in *VegfR2^RSC–/–^* kidneys (**Figure 5C**). *Tek* encodes Tie2, the receptor for ANGPT1, which regulates endothelial survival and vascular stability(17). *Klf4* encodes a transcription factor that regulates endothelial inflammatory responses and VE-cadherin-mediated barrier integrity(18), whereas *Esm1* encodes endocan, an endothelial-associated proteoglycan involved in VEGF signaling, vascular remodeling, and permeability(19). Together, these findings suggest that stromal VEGFR2 deletion is associated with altered endothelial signaling and transcriptional programs that may contribute to microvascular protection after IRI. Furthermore, *Cdh5* encodes vascular endothelial cadherin (VE-cadherin), a key component of endothelial adherens junctions that maintains cell–cell adhesion and vascular integrity(20). Consistent with increased endothelial *Cdh5* mRNA levels, immunostaining revealed preserved VE-cadherin junctional staining in *VegfR2^RSC–/–^*kidneys, compared with a more scattered and fragmented staining pattern in control kidneys 7 days after renal IRI. Together, these findings indicate that stromal VEGFR2 deletion is associated with increased expression of ANGPT1–Tie2 signaling components and preservation of endothelial junctional integrity following ischemic AKI.

## Discussion

The present study reveals an unexpected role for VEGFR2 in Foxd1-lineage renal stromal cells during ischemic AKI. Although VEGFR2 is widely recognized for its vasculoprotective functions in endothelial cells, its deletion in the renal stromal lineage attenuated tubular injury, preserved renal perfusion and microvascular integrity, and reduced endothelial oxidative stress after IRI (**Figure 6**). These findings highlight the importance of the cellular context of VEGFR2 signaling in renal vascular injury.

**Figure 6.**
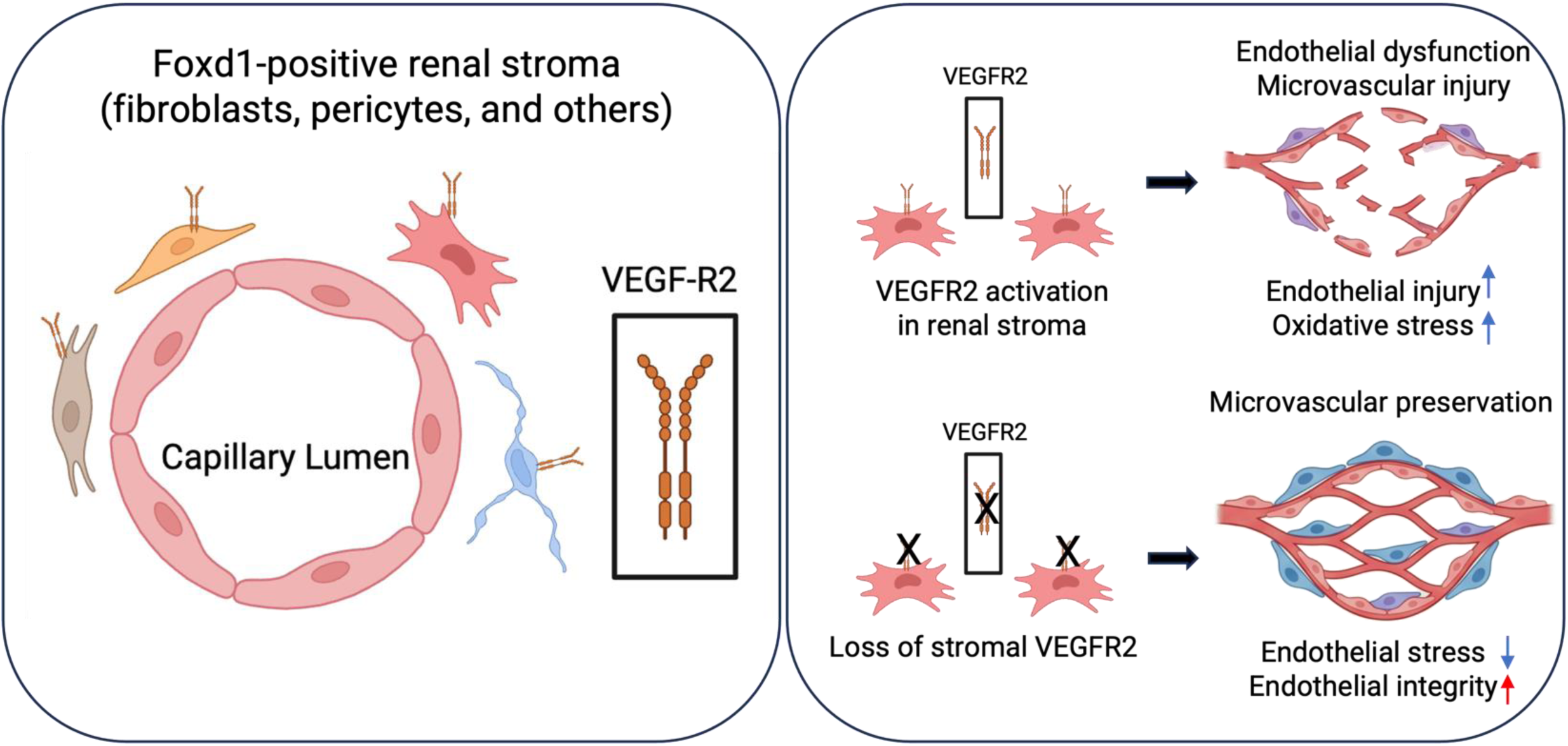
Graphical abstract: Loss of stromal VEGFR2 preserves the renal microvasculature following ischemic AKI. The figure was created using BioRender.

Renal stromal cells provide structural and paracrine support to the renal microvasculature and may regulate endothelial responses following ischemic injury(6, 7). Among Foxd1-derived stromal cell populations, pericytes closely associate with peritubular capillaries and contribute to vascular stability and endothelial homeostasis(8). Although the molecular mechanisms remain to be determined, our findings raise the possibility that altered stromal–endothelial communication contributes to the preserved microvasculature and improved renal blood flow observed in stromal *VegfR2*-deficient kidneys(9).

Although VEGFR2 has been extensively studied in endothelial cells as the principal receptor mediating VEGF-A-dependent angiogenic signaling(10), accumulating evidence suggests that VEGFR2 also functions in non-endothelial cell populations(11, 12). Developmental studies have identified a subset of Foxd1-derived stromal cells expressing endothelial markers(13). Together with our findings, these observations raise the possibility that renal stromal cells are capable of responding to VEGF signaling to support microvascular repair following ischemic injury. In the kidney, systemic VEGFR2 blockade attenuates inflammation following obstructive injury(14), but these approaches do not distinguish the contribution of individual renal cell populations. Our genetic studies therefore extend previous work by demonstrating a cell type-specific role for stromal VEGFR2 in regulating renal microvascular preservation after ischemic AKI. Recent studies have also demonstrated that VEGFR1 blockade preserves glomerular endothelial integrity in experimental diabetes, suggesting that selective modulation of VEGF receptor signaling may protect the renal microvasculature in a disease-specific context(15).

Our findings are further supported by publicly available human and mouse single-cell/nucleus transcriptomic datasets demonstrating KDR/*Kdr* expression in activated stromal cell populations following kidney injury. Moreover, independent snRNA-seq analysis of fibrotic kidneys identified *Kdr* expression in activated fibroblasts, suggesting that stromal VEGFR2 expression may represent a conserved feature of the injured kidney rather than being unique to ischemic AKI.

ANGPT1–Tie2 signaling regulates endothelial survival and vascular stabilization (14–16). We observed increased *Angpt1* and *Tek* mRNA levels, enhanced endothelial ERK phosphorylation, and more continuous VE-cadherin junctional staining in stromal *VegfR2*-deficient kidneys. These findings are consistent with an endothelial stabilization-associated response but do not demonstrate Tie2 activation or establish ANGPT1–Tie2 signaling as the cause of microvascular protection. Direct assessment of Tie2 phosphorylation and functional pathway perturbation will be needed to test this possibility.

Our findings further suggest that stromal VEGFR2 deletion is associated with an endothelial transcriptional response favoring vascular integrity following ischemic injury. Increased *Klf4* mRNA levels are particularly noteworthy, as KLF4 has been shown to transcriptionally regulate VE-cadherin and endothelial barrier function. Consistent with this, *Cdh5* mRNA levels were increased in CD31_⁺_-enriched renal cells from stromal VEGFR2-deficient kidneys, accompanied by preservation of VE-cadherin junctional staining after renal IRI. Increased *Esm1* mRNA levels further suggest altered endothelial responses to VEGF signaling and vascular remodeling, although ESM1 can also regulate vascular permeability(18, 19). Together, these observations support a model in which stromal VEGFR2 deletion influences endothelial transcriptional and junctional responses that may contribute to microvascular preservation.

An important consideration is the heterogeneity of the Foxd1 lineage, which encompasses multiple renal stromal and perivascular cell populations, including pericytes, vascular smooth muscle cells, mesangial cells, and fibroblasts. Consequently, the protective phenotype observed in *VegfR2^RSC–/–^*mice cannot be attributed to a specific stromal cell population. Future studies using more selective genetic approaches will be needed to identify the cell populations responsible for the renoprotective effects of stromal VEGFR2 deletion.

In conclusion, our findings identify an unexpected role for stromal VEGFR2 in regulating renal microvascular responses to ischemic injury. Despite the established vasculoprotective functions of endothelial VEGFR2, deletion of VEGFR2 in Foxd1-lineage renal stromal cells preserves renal perfusion, reduces endothelial oxidative stress, and maintains microvascular integrity following ischemic AKI. These findings highlight stromal–endothelial communication as a previously underappreciated determinant of renal vascular injury and suggest that selectively modulating stromal signaling may offer a novel strategy to protect the renal microvasculature. Defining the stromal cell populations and signaling mechanisms responsible for this protection may uncover new therapeutic opportunities to improve recovery from ischemic AKI.

## Methods

### Sex as a biological variable

Male mice were used throughout the manuscript. Both male and female mice were studied in the renal IRI model in Figure 2 to evaluate the effects of stromal VEGFR2 deletion in both sexes.

### sc/snRNA-Seq data analysis

scRNA-seq and snRNA-seq data for human and mouse kidneys were obtained from publicly available resources. Human kidney KDR (VEGFR2) expression was analyzed using the Kidney Precision Medicine Project (KPMP) Kidney Tissue Atlas Explorer (https://atlas.kpmp.org/explorer/dataviz). Mouse *Kdr* (*VegfR2*) expression in unilateral ureteral obstruction (UUO) day 14 kidneys was analyzed using the Kidney Interactive Transcriptomics (KIT) website developed by the Humphreys laboratory (http://humphreyslab.com/SingleCell/) (16, 17).

### Animals

To generate *VegfR2^RSC–/–^* mice, we used the transgenic *Foxd1-EGFP-Cre* mouse line, which expresses EGFP and Cre recombinase in the renal stroma(18). These mice were crossed with *VegfR2*-floxed mice (*VegfR2^fl/fl^*) to generate *Foxd1^Cre/+^; VegfR2^fl/fl^* mice, resulting in conditional deletion of *VegfR2* in the Foxd1-positive renal stroma. *VegfR2^RSC–/–^*mice have been previously characterized in our laboratory, demonstrating efficient Foxd1 lineage specific *VegfR2* deletion, and were described in Mukherjee et al(12). Age- and sex-matched Cre– littermates were used as control. All mice were maintained on an *FVB/NJ* background. Unless otherwise indicated, all experiments were performed in male mice.

### Renal IRI induced AKI model

Unilateral renal IRI followed by delayed contralateral nephrectomy was performed as previously described(30–33). Briefly, mice were anesthetized with 2% isoflurane. Core body temperature of the mice was monitored with a rectal thermometer probe and was maintained at 36.8°C–37.2°C throughout the procedures with a water-heating circulation pump system (EZ-7150; BrainTree Scientific) and an infrared heat lamp (Shat-R-Shield). Extended-release buprenorphine 1.3 mg/mL (Ethiqa XR, Fidelis MIF 900-014) was administered for analgesia (3.25 mg/kg body weight [b.w.], s.c.). With aseptic techniques, a dorsal incision was made to expose the left kidney, and renal ischemia was induced by clamping the left renal pedicle for 22 minutes in males or 31 minutes in females with a nontraumatic micro serrefine (18055–04; Fine Science Tools). Renal reperfusion was visually verified afterward. Contralateral nephrectomy of the right kidney was performed at day 6 after renal IRI. Serum and the injured left kidney were collected at day 7. Serum was analyzed by the Kansas State Veterinary Diagnostic Laboratory for a renal profile including bicarbonate, BUN, creatinine, and phosphorus.

### Real-time PCR

Real-time PCR analysis was performed as previously described to determine mRNA abundance(19). Briefly, kidneys were homogenized with a BeadBug homogenizer (Benchmark Scientific), and total RNA was isolated using the miRNeasy Mini or Micro Kit (Qiagen). cDNA was reverse transcribed from 500 ng of total RNA with SuperScript First-Strand Synthesis System II (Thermo Fisher Scientific). Real-time PCR analysis was performed with gene specific primer oligos, SsoAdvanced SYBR Green Super-mix (Bio-Rad, 1725271) and CFX96 Touch Real-Time PCR Detection System with C1000 Thermal Cycler (Bio-Rad). Cycling conditions were 95°C for 10 minutes, followed by 40 cycles of 95°C for 15 seconds and 60°C for 1 minute. Data were normalized to Rn18S as an endogenous control and analyzed using the 2^−ΔΔCT^ method(20). Each primer sequence is shown in Supplemental Table 1.

### Endothelial cells isolation

CD31 -enriched renal cells were obtained from single-cell suspensions of kidneys obtained from 10- to 14-week-old male *VegfR2^RSC−/−^* mice or Cre-negative littermate controls, 7 days after renal IRI. Cells were enriched using the Dynabeads Antibody Coupling Kit (14311D; Thermo Fisher Scientific) coupled to an anti-CD31 antibody (clone MEC 13.3, 553370; BD Biosciences), according to the manufacturer’s instructions and as previously described(34).

### Tissue section analysis

Kidneys were fixed in 4% paraformaldehyde and embedded in OCT or paraffin. These tissues were sectioned at 10 μm or 4 μm, respectively. The kidney sections were stained with H&E. H&E-stained slides were scored semi-quantitatively (from 0 to 4) for tubular damage in a blinded fashion with respect to tubular dilation, proteinaceous cast formation, and loss of brush border.

Immunostaining was performed as previously described(19), with OCT- or paraffin-embedded tissues and with following primary antibodies: anti-ANGPT1 (ThermoFisher, 23302-1-AP), anti-CD31 (ThermoFIsher, A18045), anti-Endomucin (Santa Cruz, sc-65495), anti-KIM1 (R&D Systems, MAB1817), anti-NGAL (R&D Systems, AF1857), anti-PAI-1 (ThermoFisher, MA5-17171), anti-PDGF-Rβ (ThermoFisher 14-1402-82 or MA5-15143), anti-phospho-ERK (Cell Signaling, 9101), anti-VE-Cadherin (eBioBV13, eBioscience), anti-phospho-VEGFR2 (Tyr1175) (19A10, Cell Signaling, 2478), 4HNE (ThermoFisher, MA5-27570) or LTL (Vector Laboratories, FL-1321; 1:100). Antibodies were used at 1:50–1:200, followed by conjugation with Alexa Fluor secondary antibodies at 1:200. Images were obtained with Leica DMi8 microscope equipped with Leica K5 and K3C cameras and LAS X software (Leica, Buffalo Grove, IL, USA) or Leica Stellaris 5 confocal microscope with silicon HyD detectors (Leica).

### ASL-MRI

Cortex areas of the left kidneys in mice were longitudinally analyzed by *in vivo* magnetic resonance imaging (MRI) noninvasively for anatomic and functional evaluation before and after renal IRI as reported previously(21, 22). T2-weighted MRI images were used for anatomic images. RBF rate was quantified by ASL-MRI. ASL uses the endogenous water molecules in the blood as tracers to quantify RBF without needing to administer exogenous contrast agents. A continuous adiabatic pulse train in a 1-mm slab localized 9 mm proximal to the renal artery was applied to the abdominal aorta to spin tag all of the incoming proton spins through the abdominal aorta to reach the steady state. The spin attenuation by the steady-state labeling, labeling efficiency, blood-tissue partition coefficient, and T1obs were measured to calculate the RBF with the following equation:

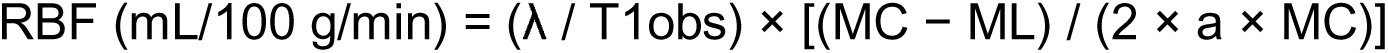

where:

- RBF = renal blood flow (mL/100 g/min)
- λ = blood–tissue partition coefficient
- T_1obs_ = observed (apparent) *T*_1_
- M_C_ = control magnetization
- M_L_ = labeling magnetization
- a = labeling efficiency

α was measured in the abdominal aorta 0.9 cm from the arterial spin inversion site. Anatomic MRI images were referred to define cortex area of the kidneys in ASL MRI images.

### Statistics

Data are presented as mean ± SD. Prism 9.0.0 software (GraphPad) was used for statistical analysis. To determine whether sample data has been drawn from a normally distributed population, D’Agostino-Pearson omnibus test or Shapiro-Wilk test was performed. For parametric data, 1-way ANOVA with *post hoc* Tukey comparison was used for multiple-group comparison and 2-tailed Student’s t test was used to compare 2 different groups. For nonparametric data, Mann-Whitney U test was used. Longitudinal renal blood flow measurements obtained by ASL-MRI were analyzed using a mixed-effects model to assess the effects of genotype, time, and their interaction, followed by *post hoc* multiple-comparison tests to evaluate differences between genotypes at individual time points. A P value less than 0.05 was considered significant.

### Study approval

All animal protocols were approved by the University of Pittsburgh IACUC (protocol no. 23103811) at the Office of Research Protection/Animal Research Protection in Pittsburgh, Pennsylvania, USA. All experiments were conducted in accordance with the guidelines and regulations of the Animal Welfare Act (AWA) and the PHS Policy on Humane Care and Use of Laboratory Animals.

## Author contributions

AO, AK, KK, AB, YS, DMC, EM, KEP, YH, KVM, NMR performed experiments and provided intellectual input. TC, JH and SSL designed and performed experiments, wrote and edited the manuscript, provided intellectual input, and oversaw the project. All authors approved the final version of this manuscript.

## Acknowledgments

This work was supported by NIDDK K01-DK133635; UPMC VMI P3HVB award; Samuel and Emma Winters Foundation Award; UPMC CHP RAC Start-up grant; UPMC CHP Foundation Children’s Trust Grantmakers’ award; George M. O’Brien Kidney Resource Alliance Opportunity Pool Program (OKRA OPP) (to TC). NIDDK R01-DK121758, DK134346, R25-DK119180 (to SSL). NIDDK R01 DK137410 (to JH). Carolyn and Mark Snyder and the Snyder Family Foundation (to JH and SSL).

The University of Pittsburgh Center for Biologic Imaging provided microscopy support. We thank the support of the UPMC CHP Animal Imaging Core Laboratory for MRI analysis, UPMC CHP Histology Core Laboratory for histology assistance, UPMC CHP Cell Imaging Core Laboratory for confocal microscopy resources, the Vanderbilt Eicosanoid Core Laboratory for tissue isofuran measurement, and the Kansas State University Veterinary Diagnostic Laboratories for serum chemistry.

---

Supplemental Figure 1. Serological analysis of *VegfR2^RSC^* ^ー^*^/^* ^ー^ kidneys demonstrates protective features in selected biomarkers after ischemic AKI. Male mouse studies. Serum analyses for phosphorus, Na/K ratio, and bicarbonate indicate that the level of renal functional impairment is decreased in *VegfR2^RSC^*^ー^*^/^*^ー^ mice 7 days after renal IRI. *p < 0.05; **p < 0.01; using 1-way ANOVA post hoc *Tukey* multiple comparison.

**Supplementary Figure 2. Stromal VEGFR2 deletion decreased immunostained PAI-1 in PDGFR**β**-positive renal interstitial cells.** PDGFRb was co-stained as an stromal derived cell marker. Scale bars: 50 μm.

**Supplementary Figure 3. Stromal VEGFR2 Deletion Upregulates Genes Associated with Vascular Integrity.** *VegfR2^RSC^*^ー^*^/^*^ー^ whole kidneys increased mRNA of vascular integrity associated genes, *Tek*, *Esm1, Proc,* and *Sdc2,* 7 days after renal IRI. *P < 0.05; **P < 0.01 using *t* test.

