## Supplemental Figures for "Loss of stromal VEGFR2 protects against ischemic acute kidney injury by preserving the renal microvasculature"

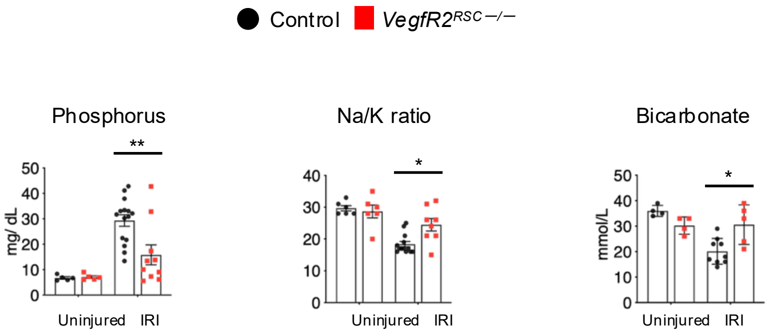


**Supplemental Figure 1.** Serological analysis of *VegfR2*$^{RSC-/-}$ kidneys demonstrates protective features in selected biomarkers after ischemic AKI. Male mouse studies. Serum analyses for phosphorus, Na/K ratio, and bicarbonate indicate that the level of renal functional impairment is decreased in *VegfR2*$^{RSC-/-}$ mice 7 days after renal IRI. *$p < 0.05$; **$p < 0.01$; using 1-way ANOVA post hoc *Tukey* multiple comparison.

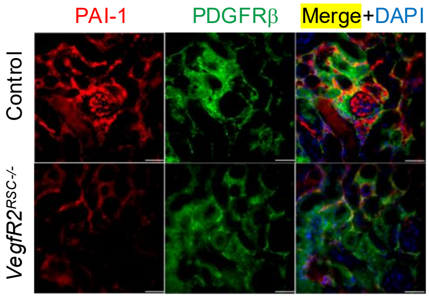


**Supplementary Figure 2. Stromal VEGFR2 deletion decreased immunostained PAI-1 in PDGFRβ-positive renal interstitial cells.** PDGFRβ was co-stained as an stromal derived cell marker. Scale bars: 50 μm.

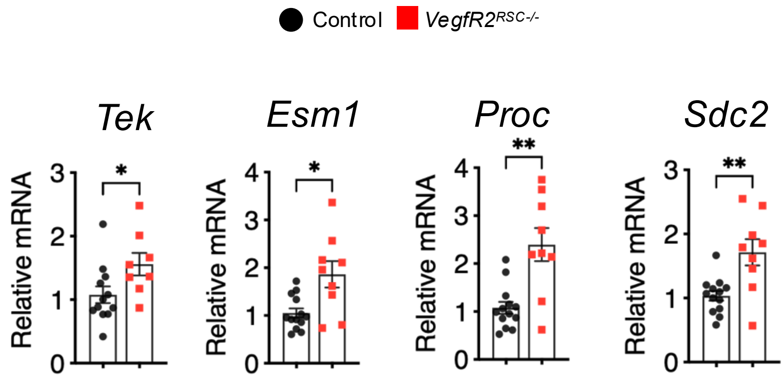


**Supplementary Figure 3. Stromal VEGFR2 Deletion Upregulates Genes Associated with Vascular Integrity.** *VegfR2*$^{RSC-/-}$ whole kidneys increased mRNA of vascular integrity associated genes, *Tek*, *Esm1, Proc,* and *Sdc2,* 7 days after renal IRI. *P < 0.05; **P < 0.01 using *t* test.
